# Cell jamming transition is regulated by mitochondrial pyruvate transport and endocytosis

**DOI:** 10.64898/2026.02.09.704880

**Authors:** Alexandra Bermudez, Zoe Latham, Johnny Diaz, Weihong Yan, Jerry Chen, Dapeng Bi, Andrew Goldstein, Jimmy K Hu, Neil Y.C. Lin

## Abstract

Epithelial tissues undergo dynamic transitions between fluid-like collective motion and mechanically jammed states during development, injury repair, and disease progression. However, the cellular programs that drive these transitions and regulate collective behavior remain unclear. Using a controlled crowding model integrated with live-cell imaging and time-resolved multi-omics, we demonstrate that epithelial crowding triggers early metabolic changes characterized by increased mitochondrial pyruvate anaplerosis that precedes the jamming transition. Functional inhibition of mitochondrial pyruvate import is sufficient to sustain collective cell motility, impeding jamming transition in crowded cells. This unjammed state is driven by enhanced cytoskeletal remodeling and requires RhoA-myosin II activity. Mechanistically, we show that elevated cytoskeletal signaling promotes macropinocytic uptake, which serves as a required feedback loop to maintain motility. These findings identify mitochondrial pyruvate utilization as a key regulator that links metabolic remodeling to the endocytic control of epithelial fluidity.

## Introduction

Epithelial tissues possess the remarkable ability to transition between fluid-like, motile collectives and solid-like, mechanically stable sheets. This ability enables the dynamic morphological and structural transformations essential for organ morphogenesis, injury repair, and tissue homeostasis^1–5^. As cells multiply within confined environments, increasing density drives extensive cytoskeletal remodeling and junction maturation. These changes trigger the contact inhibition of both locomotion and proliferation^6,7^, which subsequently reduces cell motility and rearrangements to solidify the epithelium and preserve tissue architecture^8–10^. This jamming transition is critical for stabilizing tissues during key developmental events, such as the conclusion of *Drosophila* wing elongation^11^, tissue convergent extension^12^, and the final phases of wound healing^13^. However, despite the well-characterized physical principles governing these states, the cell-intrinsic regulatory mechanisms and biological pathways that drive the jamming–unjamming transition remain a fundamental open question.

By integrating principles from mechanobiology and soft-matter physics, recent research has established a robust biophysical framework to characterize collective cell dynamics during jamming–unjamming transitions. This framework highlights the role of tissue-scale mechanics, specifically how cell packing, force transmission, and geometric deformation govern transition states^14–17^. Beyond these intrinsic mechanics, collective migration is modulated by a broad spectrum of extrinsic regulators, ranging from physical inputs such as substrate properties^18–21^ and compressive stress^22^ to environmental stimuli including morphogen gradients^23^, electrotaxis^24^, and interstitial flow^25,26^. Crucially, in many contexts, collective migration is facilitated not by an epithelial–mesenchymal transition (EMT), but by a physical unjamming transition (UJT)^22,27–30^. This UJT allows tissues to adopt a fluid, motile state while strictly maintaining their epithelial molecular identity and barrier integrity.

While the physical principles of epithelial jamming are well characterized, the cell-intrinsic pathways that transduce crowding cues into behavioral shifts remain incompletely defined^4,10^. Cadherin-mediated adhesion is a known rheological rheostat for the fluid-to-solid transition^30^; however, because upstream signals dictate adhesion dynamics^31^, the primary sensing mechanisms governing unjamming remain elusive. Elucidating these pathways is essential, as morphogenesis, wound healing, and malignancy all depend on mechanical state changes regulated by diverse cellular programs, including metabolism^32,33^.For instance, the fluidized state in wound repair requires elevated glycolytic activity^34^, whereas metabolic reprogramming during cancer invasion scales with the mechanical properties of the tumor microenvironment^35–38^. Despite these correlations, the precise mechanisms by which intracellular metabolism integrates crowding-induced mechanical information to drive the jamming transition remain an unresolved frontier.

Here, we address this question using a controlled epithelial crowding model based on Madin–Darby canine kidney (MDCK) monolayers that progress from sparse, motile collectives to densely packed, jammed sheets. By integrating live-cell imaging with time-resolved transcriptomics and metabolomics, we captured the evolving relationship between cell behavior, transcriptional remodeling, and metabolic state during the jamming transition. This multimodal framework allows us to connect molecular regulation to emergent collective dynamics and helps bridge a long-standing disconnect between physical and biological views of epithelial jamming. Using this cross-discipline approach, we identify pyruvate anaplerosis as a central response to crowding that modulates actomyosin activity and suppresses endocytosis, thereby promoting the cessation of cell movement and the establishment of a jammed state. This intrinsic program coordinates metabolic and cytoskeletal responses to increasing cell density and provides a foundation for understanding how epithelial tissues autonomously regulate transitions between fluid and solid states.

## Results

### Early Metabolic Remodeling Precedes Adhesion and YAP Translocation During Cell Crowding

Cell crowding is a dynamic process in which coordinated changes in local cell density, junction organization, and molecular signaling progressively remodel tissue mechanics. While understanding how these changes evolve is essential for elucidating the fluid-to-solid jamming transition, a detailed temporal description of these events remains lacking. To capture this evolution, we employed an MDCK epithelial monolayer model, tracking cells from a sparse, sub-confluent state to a densely packed, jammed configuration. Across fifteen evenly spaced time points (T1–T15), we examined three key processes linked to jamming: cell movement, E-cadherin–based adhesion, and YAP localization (Fig. 1A). We quantified the root-mean-square cell velocity (*V*_*rms*_) via particle image velocimetry, observing a characteristic decay and plateau that confirmed our time series captured the full jamming process (Fig. 1B). Simultaneously, we monitored the maturation of E-cadherin junctions through the immunofluorescence junctional contrast normalized to the cytoplasmic background (Fig. 1C, E, and Extended Figure 1A) and tracked the mechanoresponsive inactivation of YAP via its cytoplasmic-to-nuclear intensity ratio (Fig. 1D, F).

**Figure 1.**
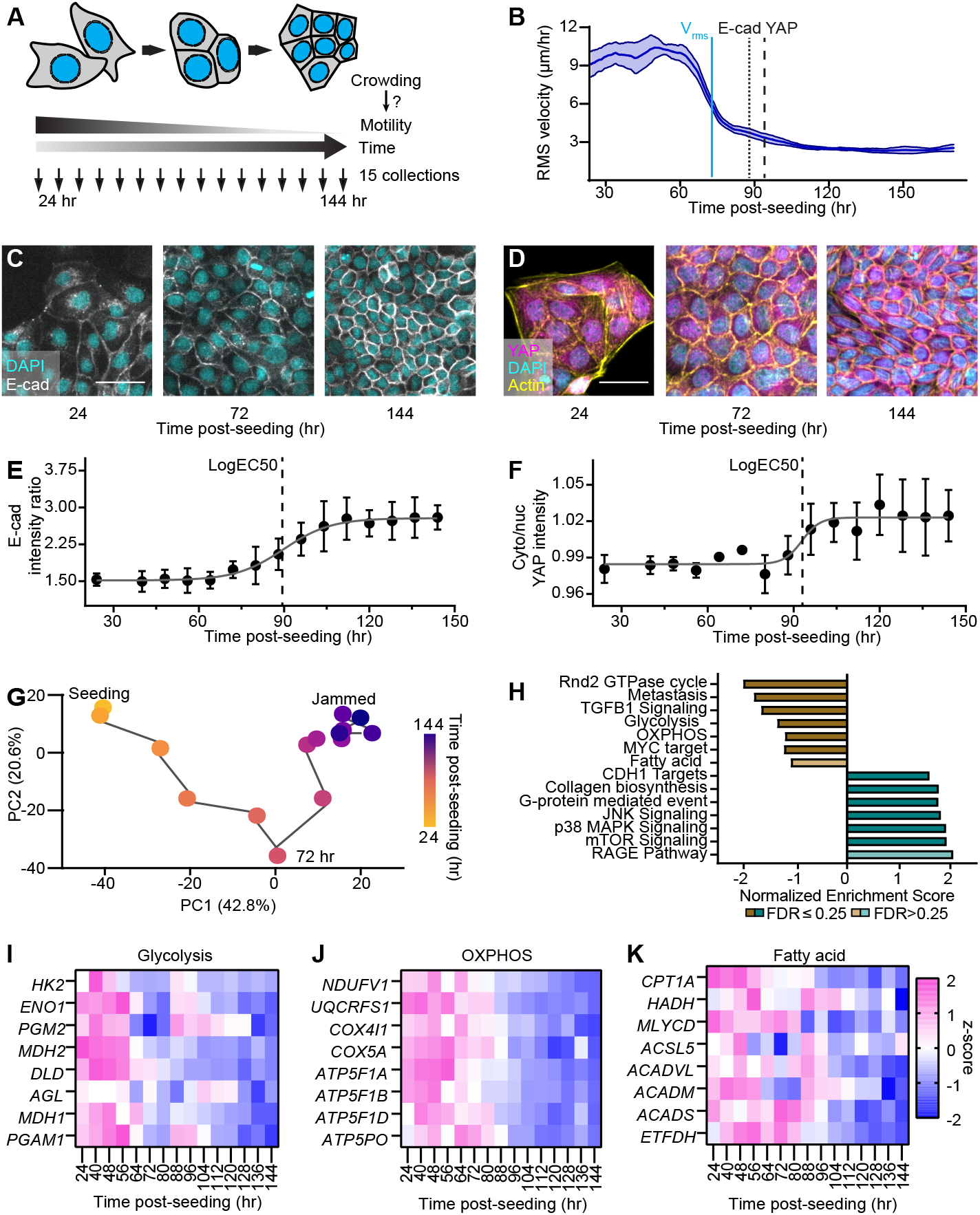
Metabolic Remodeling Precedes Adhesion and YAP Transitions in Crowding Epithelia. (A) Schematic of the experimental design using the MDCK cell jamming model. Subconfluent cells progressively undergo crowding, during which cell velocity, E-cadherin and YAP levels, and RNA expression were quantified across 15 time points. (B) Root-mean-square cell velocity (*v*_rms_) measured from the same time series exhibits an earlier decline (≈70 h) than the E-cadherin and YAP transitions. (C) E-cadherin immunofluorescence showing the progressive maturation of adherens junctions during crowding. Scale bar = 50 *µ*m. (D) YAP immunofluorescence demonstrating the gradual enrichment of YAP in the cytoplasm. (E) Quantification of the junction-to-cytoplasm E-cadherin intensity ratio reveals a time-dependent increase following a sigmoidal trend (LogEC_50_ ≈ 90 h). Scale bar = 50 *µ*m. (F) Quantification of the cytoplasmic-to-nuclear YAP intensity ratio shows a sigmoidal increase (LogEC_50_ ≈ 90 h), indicating reduced nuclear YAP localization. (G) Principal component analysis of bulk RNA-seq profiles depicts a nonlinear transcriptional trajectory with an inflection near 70 h. (H) Gene set enrichment analysis identifies pathways associated with epithelial differentiation, signaling, and metabolism that significantly change over time. (I–K) Expression dynamics of key metabolic gene sets, glycolysis, OXPHOS, and fatty-acid metabolism, show coordinated downregulation beginning near 70 h, preceding adherens junction maturation and YAP relocalization.

Notably, we found that both the initiation of E-cadherin junction maturation and the decrease in nuclear YAP occurred at approximately 90 h post-seeding, whereas cell motility (*V*_*rms*_) began to decline roughly 30 h before these events (Fig. 1B). This temporal sequence, where motility loss precedes junctional maturation and YAP relocalization, is consistent with previous studies^39,40^ showing that epithelial cells lose motility at intermediate densities well before reaching high-density confluent states. Collectively, these findings suggest that the initial regulation of collective cell motility during the jamming transition involves mechanisms independent of adherens junction maturation and YAP-dependent transcriptional control.

To identify the molecular mechanisms governing this early decline in collective motility, we performed time-series bulk RNA sequencing (RNA-seq) across the fifteen time points previously described (T1–T15), with three biological replicates per point. Principal component analysis (PCA) revealed a nonlinear transcriptional trajectory with a distinct inflection point along the second principal component (PC2) at approximately 70 h. This shift coincided with the onset of E-cadherin junction maturation and the reduction in nuclear YAP localization (Fig. 1G). Crucially, we observed pronounced and sustained transcriptional changes during the early crowding process (T1–T5) that preceded this 70 h inflection point (Fig. 1H).

Gene set enrichment analysis identified significant remodeling of cellular metabolism, specifically within glycolysis (Fig. 1I and Extended Data Fig. 1B), oxidative phosphorylation (OXPHOS; Fig. 1J and Extended Data Fig. 1C), and fatty acid metabolism (Fig. 1K and Extended Data Fig. 1D). Heatmap visualizations of z-scored metabolic genes revealed a consistent downregulation during this early phase of crowding, occurring well before the major E-cadherin and YAP transitions (Figs. 1I– K). Similar changes were observed in other gene sets, including TGF*β* 1 signaling and CDH1 targets (Extended Data Fig. 1E, F). Together, these patterns suggest that metabolic remodeling is not merely a consequence of jamming, but rather an early regulatory event in the crowding-induced transition.

### Glucose Oxidation Limits Collective Cell Motility During Epithelial Crowding

Following the identification of these early transcriptomic shifts, we sought to functionally characterize how cell crowding progression affects cellular energy production. We utilized a Seahorse XF Analyzer to perform real-time extracellular flux analysis, measuring the oxygen consumption rate (OCR) and extracellular acidification rate (ECAR) as indicators of mitochondrial oxidative phosphorylation and glycolytic activity, respectively (Fig. 2A).

**Figure 2.**
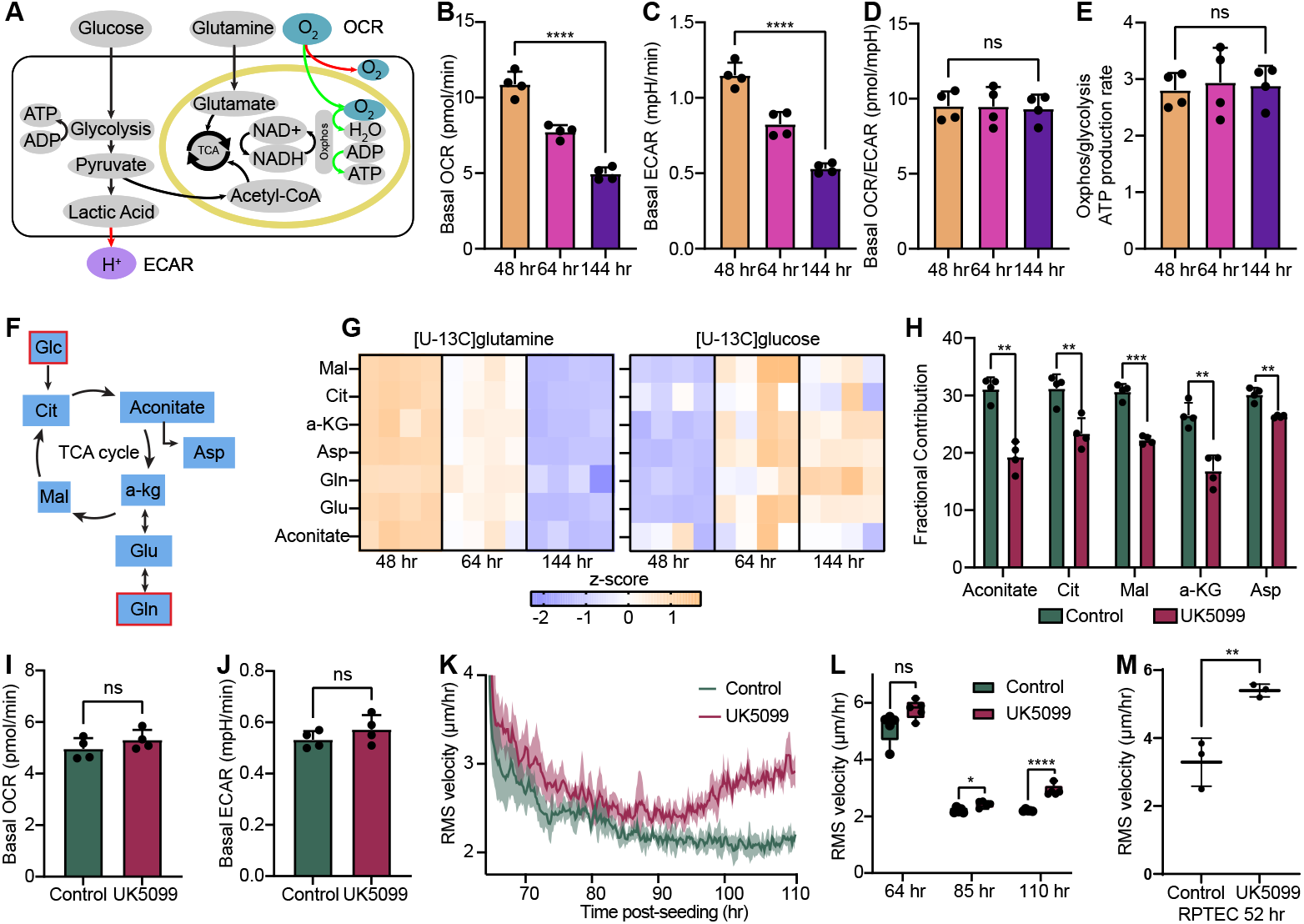
Pyruvate Oxidation Limits Motility During the Epithelial Crowding. (A) Seahorse assays were used to assess bioenergetic changes during crowding. OCR reflects oxidative phosphorylation supported largely by glucose-and glutamine-derived substrates, whereas ECAR primarily reports glycolytic proton efflux. (B–C) Both basal OCR and ECAR decrease by approximately 50% over the crowding time course. (D) The OCR/ECAR ratio remains stable, indicating a proportional downscaling of oxidative phosphorylation and glycolysis. (E) The relative ATP production from OXPHOS versus glycolysis also remains constant across time points. (F) Stable-isotope tracing with [U-^13^C]glutamine and [U-^13^C]glucose was performed to evaluate substrate routing into the TCA cycle. (G) Glutamine-derived carbon incorporation into TCA intermediates decreases over time, consistent with the global suppression of energy metabolism observed in Seahorse analyses, whereas glucose-derived carbon incorporation increases progressively. (H) Inhibition of mitochondrial pyruvate import with the MPC inhibitor UK5099 markedly reduces glucose-derived carbon labeling of TCA intermediates, confirming effective blockade of pyruvate entry. (I–J) UK5099 treatment has minimal impact on basal OCR or ECAR, indicating preserved global respiration and glycolytic activity. (K–L) MPC inhibition delays the onset of the jamming transition and significantly increases collective cell motility, as reflected by elevated *v*_rms_ during crowding. (M) This increase in *v*_rms_ upon MPC inhibition is reproduced in RPTEC monolayers, demonstrating that pyruvate-dependent mitochondrial metabolism broadly limits epithelial motility during crowding. ns, *, **, *** and **** correspond to p-values > 0.05, ≤ 0.05, ≤ 0.01, ≤ 0.001, and ≤ 0.0001, respectively.

Measurements were collected at 48 h, 64 h, and 144 h to capture the functional metabolic state during the onset of transcriptomic changes, the transitions in adhesion and YAP localization, and the attainment of a fully jammed state. We found that both basal OCR and ECAR decreased by approximately 50% over the course of the crowding process (Fig. 2B, C and Extended Data Fig. 2A), while the OCR-to-ECAR ratio (Fig. 2D) and relative ATP production rates (Fig. 2E) remained approximately stable. This coordinated decline suggests a global downscaling of cellular energy metabolism during crowding, rather than a metabolic shift between oxidative phosphorylation and glycolysis. Consequently, overall energy production decreases as cells reach a jammed state, indicating a significant reduction in metabolic activity under high-density conditions. To further clarify the metabolic underpinnings of the energy downscaling observed in our Seahorse assays, we investigated how nutrient utilization is redistributed across major metabolic pathways. We performed stable-isotope tracing using uniformly labeled [U-^13^C]glutamine and [U-^13^C]glucose, the primary carbon sources fueling the tricarboxylic acid (TCA) cycle (Fig. 2F). Metabolomic analysis (Extended Data Figs. 2B, C) revealed a progressive decline in the incorporation of glutamine-derived carbon into TCA cycle intermediates (Fig. 2G and Extended Data Fig. 2D). This reduction indicates a significant decrease in glutaminolysis-dependent anaplerosis during the jamming transition, consistent with the global suppression of energy metabolism identified via extracellular flux analysis. Together, these findings suggest a metabolic shift away from glutamine oxidation as TCA cycle replenishment transitions toward a more energetically quiescent state.

In striking contrast to the decline in glutamine utilization, [U-^13^C]glucose labeling revealed a progressive increase in the fractional incorporation of glucose-derived carbon into TCA cycle intermediates over time (Fig. 2G and Extended Data Fig. 2E), despite the overall decrease in energy production. Detailed isotopologue labeling patterns for these intermediates are provided in Extended Data Figs. 2F–K. These tracer-derived data demonstrate that cells utilize proportionally more glucose as a carbon source for the TCA cycle during crowding, presenting a paradox in which glucose contribution increases while total ATP output declines.

To address the functional significance of this paradoxical increase in glucose oxidation, we inhibited mitochondrial pyruvate import using UK5099, a selective inhibitor of the mitochondrial pyruvate carrier (MPC). As expected, UK5099 treatment markedly reduced the incorporation of glucose-derived carbon into TCA cycle intermediates, confirming the effective blockade of pyruvate entry into the mitochondria (Fig. 2H and Extended Data Fig. 3A). As expected, basal OCR and ECAR remained largely unchanged, indicating that global mitochondrial respiration and glycolytic activity were preserved under these conditions (Fig. 2I, J and Extended Data Fig. 3B). These results demonstrate that mitochondrial pyruvate restriction does not suppress global respiration, but rather specifically alters the metabolic substrate routing.

**Figure 3.**
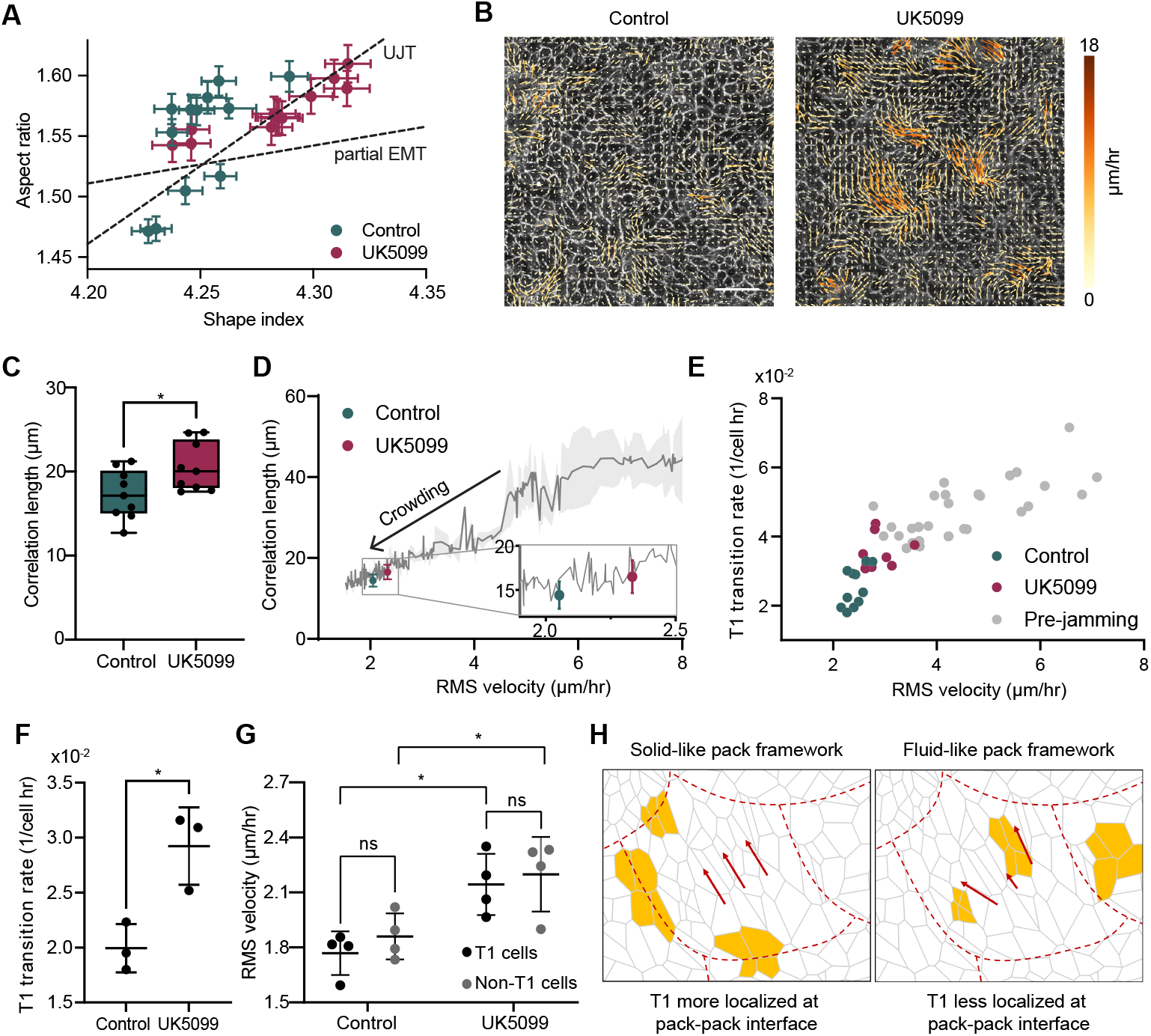
MPC inhibition induces a mechanically unjammed, fluid-like tissue state. (A) Aspect ratio and shape index distributions show UK5099-treated monolayers shift along the UJT axis (elongated, straight boundaries) rather than the partial EMT axis (tortuous boundaries), indicating fluid-like geometry with preserved epithelial integrity. (B) Representative velocity field maps from control and UK5099-treated monolayers at ∼100 h post-seeding illustrate enhanced collective motion under MPC inhibition. Scale bar = 50 *µ*m. (C) UK5099 produces an increase in the velocity correlation length *ξ*_*v*_, indicating more long-range coordination of cell movement. (D) Plotting *ξ*_*v*_ against *v*_rms_ shows that MPC inhibition induces a motility–coordination relationship characteristic of a pre-jamming, fluid-like state. The solid line represents the trajectory of untreated monolayers undergoing a jamming transition; shading denotes s.d. (E) The rate of T1 neighbor-exchange events exhibits an approximately linear dependence on *v*_rms_, consistent with increasing tissue fluidity. (F) UK5099 increases T1 transition frequency by ∼ 30–50%. (G) Instantaneous velocities of cells undergoing T1 transitions are comparable to non-T1 neighbors in both control and treated monolayers. (H) Together, these results suggest that MPC inhibition promotes coordinated yet fluid-like collective motion: correlation lengths increase with motility, while T1 events occur broadly throughout the tissue rather than being localized to pack interfaces. ns and * correspond to p-values > 0.05 and ≤ 0.05, respectively.

This inhibition of mitochondrial pyruvate import, however, led to a pronounced enhancement of collective cell motility during the crowding process (Fig. 2K, Extended Data Fig. 3C, and SI Video 1). Within 24 h of treatment (added at the 48 h time point), UK5099-treated cells displayed an approximately 40% increase in average migration speed relative to vehicle-treated controls at the 110 h mark (Fig. 2L). This effect was reproduced in human renal proximal tubule epithelial cells (RPTEC), demonstrating that pyruvate-dependent mitochondrial metabolism broadly limits epithelial motility during crowding (Fig. 2M). We also ruled out glutamine depletion as the cause of this phenotype, as the increased motility persisted in cultures supplemented with GlutaMAX (Extended Data Fig. 3D). Together, these findings demonstrate that mitochondrial pyruvate anaplerosis is critical for dampening cell motility, and inhibiting this pathway alters the metabolic state to promote migratory behavior during crowding.

### MPC Inhibition Promotes Coordinated Fluid-like Dynamics in Epithelial Monolayers

Building on the observation that UK5099 treatment enhances cell motility and delays the normal progression toward a jammed state, we examined how MPC inhibition impacts fundamental cell properties and epithelial identity. We found that MPC inhibition had negligible effects on cell proliferation and survival, as indicated by the proportion of EdU-positive and cleaved CASP3-positive cells, respectively. Furthermore, overall cell density remained unaffected by the treatment (Extended Data Fig. 4A–E).

Epithelial adhesion was also largely preserved in UK5099-treated cells, with no significant loss of total E-cadherin expression (Extended Data Fig. 4F–I). While we observed increased boundary tortuosity in treated cells (Extended Data Fig. 4I), the levels of homophilically bound E-cadherin, representing the active state of the junctional protein, showed no significant difference in mean intensity between control and UK5099-treated conditions (Extended Data Figs. 4J and K). These results indicate that the observed increase in motility occurs while maintaining the cells’ core epithelial molecular identity and barrier integrity.

The preservation of epithelial identity suggests that the observed motility increase is associated with a mechanical jamming– unjamming transition. Within the jamming framework, unjamming represents a shift from a solid-like, immobile collective to a fluid-like state without the loss of epithelial characteristics. This process is often accompanied by stereotypical changes in cell morphology, specifically increases in the cell aspect ratio and the cell shape index 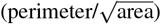 as tissues transition toward fluidization.

We, therefore, quantified these parameters (Extended Data Fig. 5A, B) and found that UK5099-treated monolayers exhibited right-shifted distributions in both metrics (Fig. 3A). These shifts correspond to more elongated cell geometries and elevated shape indices compared to controls. We found that the slope of this morphological distribution differed from the profile characteristic of partial epithelial-mesenchymal transition (EMT), where increased boundary tortuosity raises the shape index independently of the aspect ratio (Extended Data Fig. 4I). These findings indicate that MPC inhibition promotes a mechanical unjamming transition (UJT) while preserving epithelial integrity and avoiding EMT-like remodeling.

Having established a morphological signature of unjamming following MPC inhibition, we next investigated whether UK5099 also enhances the coordination of motion within the epithelial sheet (Fig. 3B). The velocity correlation length, *ξ*_*v*_, which measures the spatial persistence of velocity correlations across the tissue, serves as a quantitative indicator of collective dynamics: longer correlation lengths reflect greater mechanical coupling and coordinated movement, whereas shorter lengths characterize localized, jammed behavior^41^. UK5099 treatment produced a modest but significant increase in correlation length (Fig. 3C and Extended Data Fig. 5C), indicating more cohesive, long-range coordination among neighboring cells. Within the jamming–unjamming framework, this shift signifies a transition toward a more fluid, pre-jamming state (Fig. 3D). The emergence of this coordinated, “pack-like” motion provided the basis for our subsequent analysis of collective topological rearrangements.

To evaluate how elevated coordination in cell motility modulates tissue architecture, we quantified T1 neighbor-exchange events (Extended Data Fig. 5D and SI Videos 2 and 3), a topological metric that signals increased fluidity approaching the unjamming point. In a rigid-pack model of collective motion, T1 transitions are typically confined to interfaces between cell packs and are expected to depend only weakly on mean cell speed^41^. Using a simple scaling argument, we find that if T1 transitions occur primarily at cluster interfaces, the fraction of boundary cells scales as *f*_*int*_ ∼ 1*/L*_*c*_ ∼1*/V*_*rms*_ (where *L*_*c*_ is the cluster size). Since the relative speed between neighboring clusters scales as *V*_*rel*_ ∼ *V*_*rms*_, the total T1 rate *R*_*T*1_ ∼ *f*_*int*_*V*_*rel*_ should theoretically remain independent of *V*_*rms*_. Contrary to this expectation, UK5099-treated monolayers exhibited approximately 30% higher T1 frequencies than untreated controls under identical crowding conditions (Fig. 3E). This increase in T1 transition rate (Fig. 3F) is consistent with a fluid-like unjamming transition, further supported by the lack of global planar polarity (Extended Data Figs. 5E and F) and elevated isotropic (Extended Data Figs. 5G and H) and deviatoric (Extended Data Figs. 5I and J) strain rates. These results collectively indicate that UK5099 not only elevates collective cell movement but also promotes more frequent neighbor exchanges, reflecting an unjamming behavior distinct from simple pack translation and consistent with enhanced local fluidity across the tissue.

Finally, to link motility to topological dynamics at the single-cell level, we compared the instantaneous speeds of cells undergoing T1 transitions with those of non-rearranging neighbors. In both control and UK5099-treated conditions, T1 and non-T1 cells exhibited comparable velocities (Fig. 3G). Combined with the observed increases in velocity correlation length and T1 transition frequency, this result indicates that UK5099 promotes a partially coordinated yet fluid tissue state: cells move in fluid-like packs whose characteristic size increases with motility, while topological rearrangements remain widespread across the monolayer rather than being localized to pack interfaces (Fig. 3H). Altogether, these morphological, kinematic, and topological observations demonstrate that UK5099 induces a mechanical unjamming transition characterized by preserved epithelial cohesion, enhanced collective coordination, and increased dynamic rearrangements within the cell layer.

### MPC Inhibition Promotes Motility through RhoA-dependent Cytoskeletal Remodeling

To elucidate how MPC inhibition drives increased cell motility, we next examined alterations in actin organization, as actin remodeling directly governs cell movement. To this end, we established an MDCK cell line expressing LifeAct-GFP to visualize filamentous actin (F-actin) in live cells. Upon treatment with UK5099, we found that cells exhibited a pronounced increase in the number of stress fibers (Figs. 4A and B), while the average thickness of individual fibers remained unchanged (Extended Data Figs. 6A and B).

**Figure 4.**
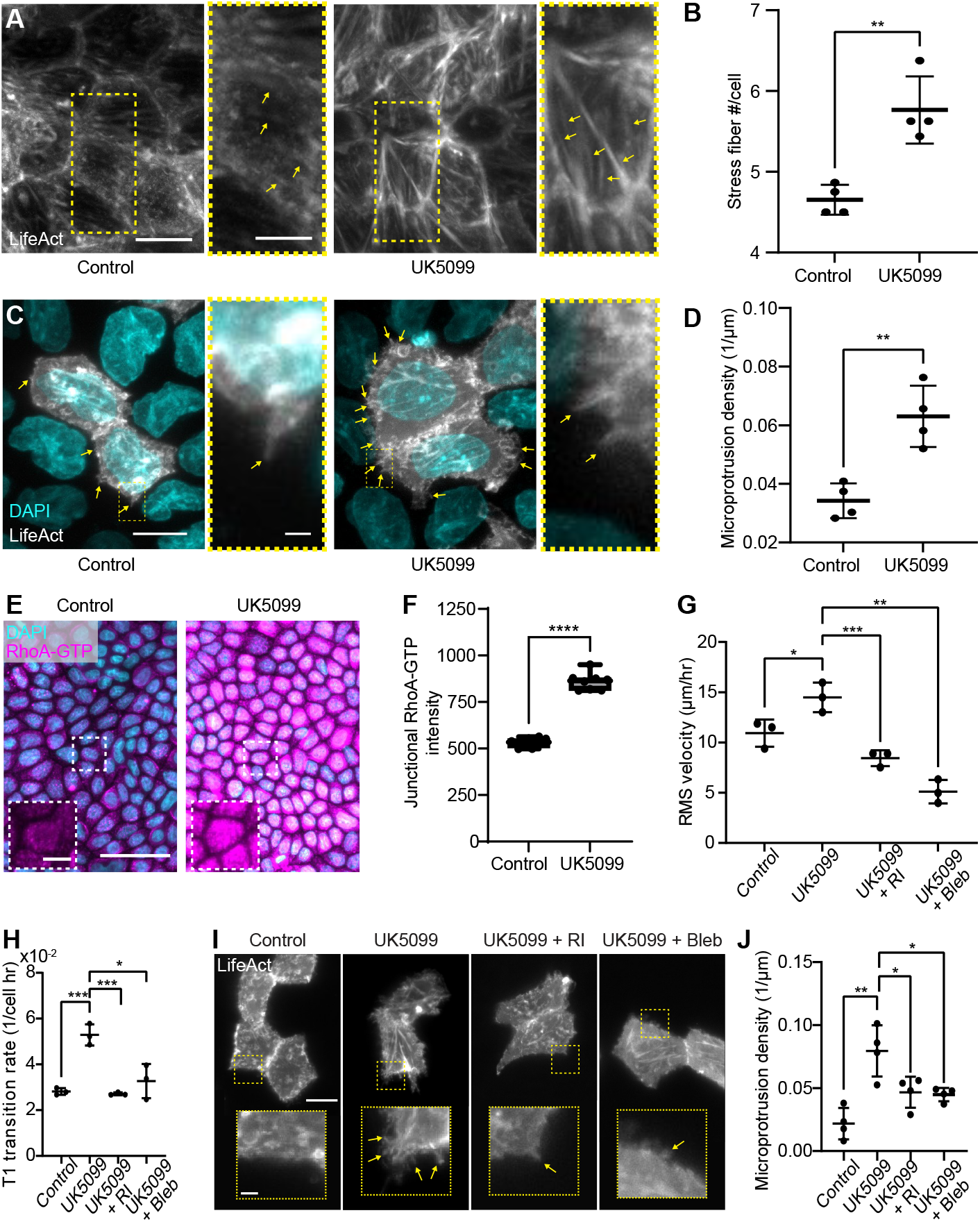
MPC inhibition enhances RhoA-dependent cytoskeletal remodeling to promote motility. (A) LifeAct-GFP imaging reveals an increase in the number of stress fibers in UK5099-treated monolayers. Scale bar = 10 *µ*m. (B) Quantification confirms elevated stress fiber density. (C) Sparse LifeAct-GFP labeling shows that MPC inhibition increases the frequency of cryptic lamellipodia (microprotrusions). Scale bars = 10 *µ*m and 1 *µ*m, respectively. (D) Microprotrusion density is significantly elevated in UK5099-treated cells. (E) Immunostaining for active RhoA-GTP demonstrates higher signal intensity following MPC inhibition, indicating increased RhoA activity. Scale bar = 50 *µ*m. (F) Junctional RhoA-GTP intensity is significantly elevated. (G) Inhibiting downstream RhoA–ROCK signaling (Y-27632) or myosin II activity (blebbistatin) abolishes the UK5099-induced increase in cell motility. (H) The UK5099-driven rise in T1 neighbor-exchange events is similarly suppressed by ROCK or myosin II inhibition. (I–J) Both ROCK and myosin II inhibition markedly reduce microprotrusion formation and density. Scale bars = 10 *µ*m and 1 *µ*m, respectively. *, **, *** and **** correspond to p-values ≤ 0.05, ≤ 0.01, ≤ 0.001, and ≤ 0.0001, respectively.

Since increased actin assembly often correlates with greater actin-driven protrusive activity, we quantified the density of cryptic lamellipodia – thin, actin-rich protrusions that extend beneath neighboring cells to mediate traction transmission during collective migration. Because these fine structures are easily obscured in confluent, uniformly labeled monolayers, we employed a sparse-labeling strategy, mixing a small fraction of LifeAct-GFP-expressing cells with unlabeled cells (Extended Data Fig. 6C). Using this system, we found that UK5099-treated cells displayed a significantly higher frequency of cryptic lamellipodia (Figs. 4C and D), though the average protrusion length remained unchanged (Extended Data Fig. 6D). These results are consistent with elevated actin turnover and enhanced coordination of collective motility following MPC inhibition.

We next investigated whether the enhanced actin remodeling observed in UK5099-treated cells was driven by elevated activity of RhoA, a key regulator of actin assembly. Immunostaining for active GTP-bound RhoA using a conformation-specific antibody revealed an approximately 60% increase in signal intensity, both within individual cells and along cell-cell junctions, when MPC function was blocked (Figs. 4E and F). This increase was accompanied by a rise in total RhoA levels (Extended Data Figs. 6E and F) and moderately elevated active Rac1-GTP intensity (Extended Data Figs. 6G and H).

To test the functional significance of RhoA in these phenotypes, we simultaneously inhibited MPC and ROCK, a crucial downstream effector of RhoA that promotes actin polymerization and myosin II activation. Treatment with the ROCK inhibitor Y-27632 significantly decreased cell motility (Fig. 4G), the frequency of T1 transitions (Fig. 4H), and the density of peripheral microprotrusions (Figs. 4I and J). Consistent with these results, inhibition of myosin II using blebbistatin similarly abolished these activities (Figs. 4G–J). Together, these findings identify RhoA and actomyosin as central mediators of the UK5099-induced increase in actin remodeling and cell motility. Mitochondrial pyruvate restriction thus enhances RhoA-ROCK-dependent actin assembly, driving cytoskeletal reorganization and motility during epithelial crowding.

### MPC Inhibition Enhances Endocytic Remodeling to Drive Epithelial Cell Motility

Live-cell imaging of LifeAct-GFP-expressing epithelial monolayers revealed that UK5099-treated cells frequently exhibited dynamic membrane ruffling followed by the formation of circular enclosed structures (SI Videos 4 and 5). These features are characteristic of macropinocytosis, an actin-driven, clathrin-independent form of endocytosis that enables the internalization of extracellular fluid and serves as an auxiliary nutrient acquisition pathway^42^. The induction of such activity upon MPC inhibition likely represents an adaptive response to altered mitochondrial metabolism, while the enhanced membrane ruffling observed under these conditions may further facilitate cell motility.

To investigate this link, we first quantified endocytic activity by immunostaining for EEA1, a canonical marker of early endosomes, and found that MPC inhibition markedly increased the density of EEA1-positive vesicles (Fig. 5A, B). This effect was fully reversed by treatment with EIPA (5-(N-ethyl-N-isopropyl) amiloride), an inhibitor of the Na^+^/H^+^ exchanger required for both macropinocytic uptake and early endosome formation. We further found that the expansion of early endosomes was reversed by Y-27632 treatment (Fig. 5C, D), suggesting that RhoA activity mediates this endocytic upregulation. In contrast, inhibiting endocytosis had negligible effects on cytoskeletal organization, such as stress fiber density (Extended Data Fig. 7A, B). Dextran uptake assays revealed enhanced bulk fluid-phase endocytosis in UK5099-treated cells (Fig. 5E, F), further supporting an overall increase in macropinocytic capacity.

**Figure 5.**
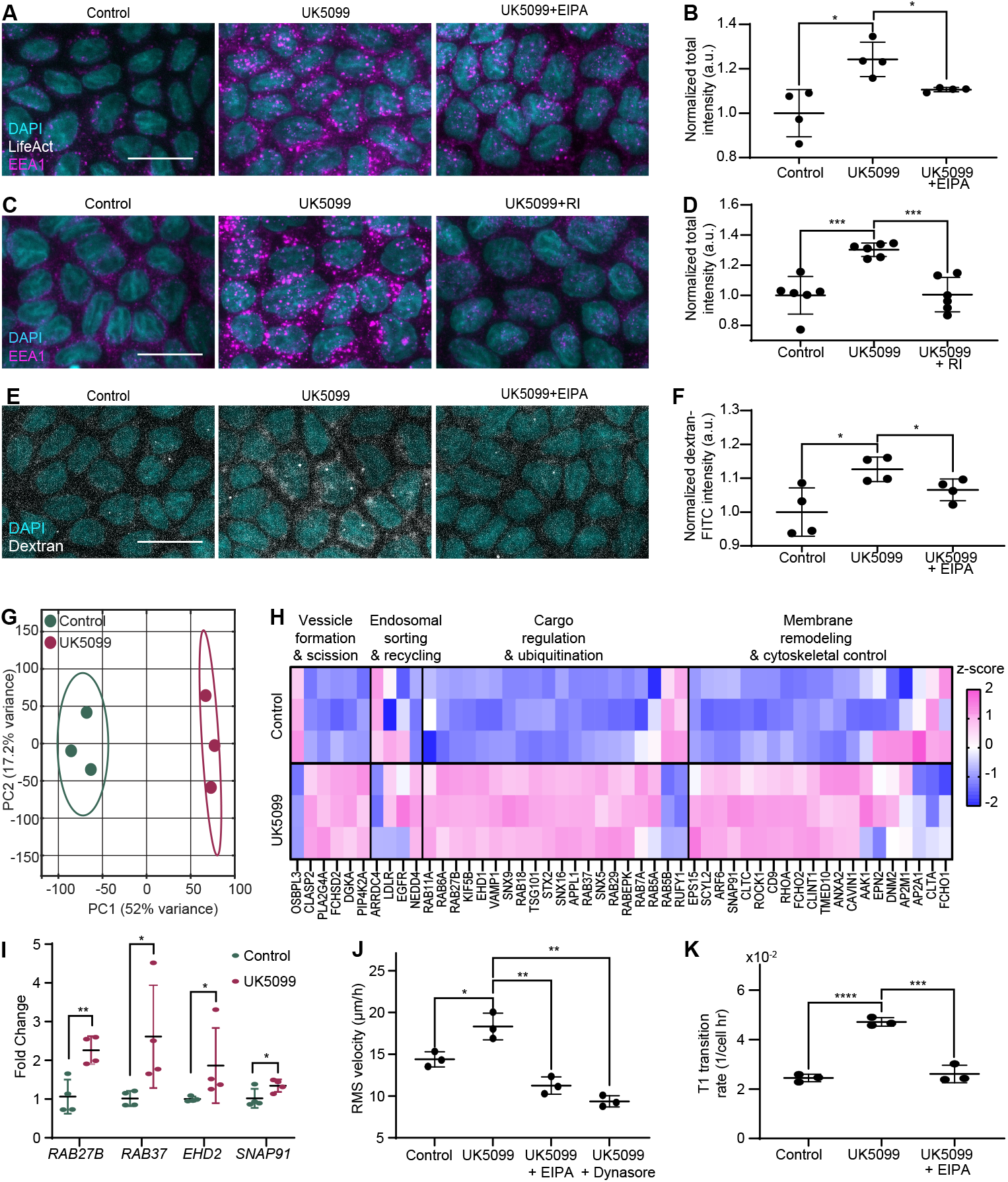
MPC inhibition enhances macropinocytosis to drive epithelial motility. (A–B) Representative immunofluorescent images (A) and quantification (B) of the early endosome marker EEA1. Treatment with UK5099 increases EEA1 intensity, which is reversed by the Na^+^/H^+^ exchange inhibitor EIPA. (C–D) Representative images (J) and quantification (K) of EEA1 staining showing that Rho-kinase inhibition (Y-27632) prevents the UK5099-induced expansion of early endosomes. Scale bar = 20 *µ*m for A, C, and E. *, **, *** and **** correspond to p-values ≤0.05, ≤0.01, ≤0.001, and ≤0.0001, respectively. (E–F) Representative images (E) and quantification (F) of Dextran-FITC uptake assays, indicating that UK5099 enhances bulk fluid-phase endocytosis (macropinocytosis) in an EIPA-sensitive manner. (G) PCA plot of RNA-seq data showing distinct transcriptional separation between control and MPC-inhibited (UK5099) samples. (H) Heatmap displaying the coordinated upregulation of major endocytosis-associated genes in UK5099-treated cells. (I) qPCR validation confirms the upregulation of key vesicular trafficking genes (e.g., *RAB27B, SNAP91*). (J–K) Functional analysis showing that blocking macropinocytosis with EIPA abolishes the UK5099-induced increase in cell motility (J) and reduces the T1 transition rate (K).

To uncover the transcriptional basis of this enhanced endocytic activity, we performed bulk RNA-seq analysis, which revealed a clear segregation between control and UK5099-treated samples in principal component space. This distinct clustering reflects global transcriptional reprogramming following metabolic perturbation (Fig. 5G). Our analysis specifically highlighted an upregulation of RhoA-associated genes (Extended Data Fig. 7C) and confirmed that EMT activation was absent (Extended Data Fig. 7D). Furthermore, analysis of endocytosis-related transcripts showed a coordinated upregulation in UK5099-treated cells, as visualized by z-scored expression heatmaps (Fig. 5H). Several key regulators of vesicular trafficking, including *RAB27B, RAB37, EHD2*, and *SNAP91*, were significantly upregulated, a result we validated via quantitative PCR (Fig. 5I). These findings indicate that MPC inhibition fundamentally upregulates the expression of the endocytic machinery.

Finally, to test whether this increased macropinocytic activity is functionally required for the observed motility phenotype, we treated cells with EIPA in the presence of UK5099. Inhibition of macropinocytosis completely abolished the UK5099-induced increase in cell motility (Fig. 5J) and the T1 transition rate (Fig. 5K), demonstrating that endocytic remodeling is essential for the migratory response.

Together, these findings establish that MPC inhibition enhances macropinocytosis, which in turn promotes collective epithelial cell motility. This discovers a signaling axis that links altered mitochondrial metabolism to the mechanical and behavioral reprogramming of epithelial tissues through endocytic control.

## Discussion

Our findings demonstrate that metabolic programs actively modulate epithelial jamming transitions, revealing a regulatory dimension that extends beyond geometric constraints and previously recognized mechanical influences. While unjamming and collective migration have been associated with increased glycolytic and oxidative metabolism^34,43^ and ATP-intensive biosynthetic demands^44,45^, the causal influence of specific metabolic pathways on cellular jamming has remained largely untested. Here, we address this gap by showing that metabolic reprogramming regulates jamming in two key ways. First, metabolic remodeling during crowding is not merely a byproduct of jamming; rather, early in the crowding process, epithelial cells adopt a pyruvate-anaplerotic program that subsequently restrains motility. Perturbing this program is sufficient to enhance cell movement and delay jamming without inducing a mesenchymal fate. Second, this regulatory mode is independent of overall energy production, as restricting mitochondrial pyruvate import alters motility despite minimal effects on oxidative phosphorylation or total ATP generation. Together, these results identify MPC-dependent pyruvate anaplerosis as a key regulator of epithelial jamming, distinct from the energy-producing glycolytic states associated with mechanically induced unjamming^34^.

We further show that disrupting mitochondrial pyruvate import induces a fluid-like epithelial state that aligns with the hallmark features of unjamming. MPC inhibition increased velocity-correlation lengths and produced coordinated, pack-like flows characteristic of fluidized epithelia^22,28,46,47^. This inhibition also elevated the frequency of T1 transitions, consistent with theories identifying T1 rearrangements as a primary route to tissue fluidization^48,49^. The comparable speeds observed between T1-engaged and non-T1-engaged cells further support a model in which fluidity emerges from collective dynamics rather than from a subset of highly motile cells within relatively rigid packs^41^.

This fluidization phenotype arises through a metabolic shift that triggers RhoA–actomyosin remodeling. Inhibiting MPC elevated active RhoA, augmented stress fiber density, and increased the formation of cryptic lamellipodia, all of which were reverted by either ROCK or myosin II inhibition. These findings indicate that mitochondrial pyruvate metabolism directly modulates actomyosin organization, consistent with reports linking metabolic states to RhoA and cytoskeletal activity. For instance, previous studies have shown that oxidative and nitrative stress can activate RhoA^50^, condensates of glycolytic enzymes can promote actin assembly^51,52^, and glycolytic activity can enhance lamellipodial dynamics^53^. Our findings converge with these observations to describe a model in which glucose metabolism interfaces with RhoA to drive the cytoskeletal changes required for the jamming transition.

Finally, our results demonstrate that increased endocytosis acts downstream of the MPC-RhoA–actomyosin axis to sustain cell motility in crowded epithelia. This echoes prior work showing that enhanced endocytosis promotes epithelial fluidization^27,54,55^; in particular, RAB5A-induced early endosome formation and macropinocytosis are known to unjam crowded epithelia^28^. Our study reveals mitochondrial pyruvate utilization as a key upstream regulator of this endocytosis-driven unjamming response. This discovery establishes a signaling axis linking cellular crowding to pyruvate oxidation, ROCK-dependent actin remodeling, and the endocytic control of epithelial mechanics. Ultimately, this work unveils a previously unappreciated metabolic mechanism through which epithelial cells regulate endocytic activity to control the jamming-–unjamming transition during tissue crowding.

## Methods

### Cell Culture and Drug Treatment

MDCK cells were cultured in Minimum Essential Medium Alpha (MEM-*α*; Fisher Scientific, 12561-056) supplemented with 10% fetal bovine serum (FBS; Fisher Scientific, 12662-029) and 1% Penicillin–Streptomycin (Fisher Scientific, 15140-122). Cells were maintained at 37 °C in a humidified incubator with 5% CO_2_. For routine maintenance, cells were passaged at approximately 80% confluence using Trypsin–EDTA solution (Fisher Scientific, 25300-054). Experimental cultures were seeded at a density of 30,000 cells/cm^2^. For inhibition of the mitochondrial pyruvate carrier (MPC), cells were treated with 2 or 5 *µ*M UK5099 (Selleckchem, S5317). These concentrations were optimized to achieve effective MPC inhibition while minimizing cytotoxicity or induction of apoptosis. Pharmacological treatments were performed at 37 °C and 5% CO_2_ near the onset of confluence, approximately 64 hours after seeding at 30,000 cells/cm^2^.

### Generation of Stable Cell Lines

To visualize F-actin dynamics, we generated a stable MDCK cell line expressing LifeAct-14–eGFP. Lentiviral particles were produced in HEK293T cells using the LifeAct-14 transfer plasmid (Addgene, 158750) and subsequently used to transduce parental MDCK cells. Following transduction, cells were selected with blasticidin to establish a homogeneous population with stable LifeAct-14–eGFP expression. Stable integrants were expanded and validated via fluorescence microscopy, which confirmed robust and consistent labeling of cortical F-actin. For experiments requiring the visualization of sub-cellular structures in confluent monolayers, these LifeAct-GFP-expressing cells were mixed with unlabeled parental MDCK cells at a low ratio to achieve sparse labeling.

### RNA-seq Analysis

Total RNA was extracted from cells using the Zymo RNA Extraction Kit (R2072) according to the manufacturer’s protocol. RNA sequencing was performed on a single lane of an Illumina NovaSeq S4 flow cell (200-cycle kit), achieving an average read depth of approximately 44 million reads per sample. Raw RNA-seq reads were first subjected to quality assessment using FastQC. Adapter sequences and low-quality bases were trimmed with Cutadapt^56^. The cleaned reads were then aligned to the domestic dog reference genome (CanFam4) using STAR (version 2.7.10a), an ultrafast RNA-seq aligner^57^. Gene annotations were derived from the NCBI RefSeq All dataset, obtained from the UCSC Genome Browser (https://genome.ucsc.edu). Gene-level read counts generated by STAR were normalized and analyzed for differential expression using the `limma` package in R^58^.

### Immunostaining

Cells were cultured in 4-chamber Ibidi slides (Ibidi, 80426) and fixed with 10% neutral-buffered formalin containing 0.03% Eosin (Sigma-Aldrich, F5304-4L) for 10 minutes at room temperature. Permeabilization and blocking were performed for 30 minutes in PBS containing calcium and magnesium (Fisher Scientific, 14040-133), supplemented with 2% donkey serum (Fisher Scientific, D9663-10ML) and 0.25% Triton X-100. Following three washes with PBS, cells were incubated for 30 minutes with the following primary antibodies: E-cadherin (Abcam, ab11512), YAP (Novus Biologicals, NB110-58358), RhoA (Cell Signaling Technology, 2117T), Rac-1 GTP (NewEast Biosciences, 26903), or EEA1 (BD Biosciences, 610456).After washing three times with PBS, cells were incubated for 30 minutes at room temperature with secondary antibodies—anti-mouse Alexa Fluor 488 or anti-rabbit Alexa Fluor 647 (Invitrogen, A21042 and 4414S, respectively)—together with DAPI (Abcam, ab228549) and either phalloidin (Invitrogen, A12379) or actin stain (Invitrogen, R37110).

### Microscopy and Live-cell Imaging

Fluorescence images were acquired using either confocal or widefield microscopy systems. Confocal imaging was performed using a Rescan Confocal Microscope (RCM1) integrated with a Nikon Eclipse Ti-E (NIS-Elements software) or an NL5+ system mounted on a Nikon Ni-E or Zeiss AXIO Observer Z1 (Micro-Manager software). Widefield imaging was conducted via an Etaluma LS720 (LumaView 720/600-Series software) or an ECHO REVOLUTION microscope.

For confocal acquisitions, we utilized 20 × WI (0.95 NA), 20 × (0.75 NA), 40 × oil (1.30 NA), and 63 × oil (1.40 NA) objectives. Widefield imaging employed a 20 × (0.40 NA) objective on the Etaluma system and a 20 × (0.75 NA) objective on the ECHO REVOLUTION microscope. Imaging parameters, including laser power and exposure times, were maintained consistently across all experimental conditions to ensure comparability. For intensity quantifications, z-projected images were generated from z-stack acquisitions with a 1.5 *µ*m step size.

### Time-lapse Collective Motility Analysis

To characterize collective cell dynamics, time-lapse imaging was performed under controlled environmental conditions. For cell motility quantification, MDCK monolayers were maintained in a humidified incubator at 37^°^C and 5% CO_2_ and imaged using the Etaluma LS720. Phase-contrast images were acquired at 15-minute intervals for up to 144 h (6 days) using a 20× Long Working Distance (LWD) U Plan Fluorite objective (NA=0.5, WD=2.1 mm, CGT=0.17 mm; Olympus UPLFLN20X). Culture media were refreshed daily during long-term acquisitions. High-resolution characterization of cytoskeletal dynamics in LifeAct-GFP-expressing cells was performed on the NL5+ confocal system with acquisition intervals ranging from 1 to 15 minutes.

### Quantification of Kinematics and Topological Rearrangements

Cellular velocity fields were quantified using Particle Image Velocimetry (PIV) analysis in `MATLAB`. The root-mean-square cell velocity (*V*_*rms*_) was calculated as: 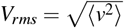 where *v* represents the instantaneous velocity vector and ⟨… ⟩ denotes the spatial average over the field of view.

Topological rearrangements were evaluated by identifying T1 neighbor-exchange events using the Tissue Analyzer plugin in `ImageJ/Fiji`. A T1 transition was defined as the collapse of a cell–cell junction shared by two cells, followed by the formation of a new junction between two previously non-neighboring cells. The T1 transition rate was calculated as the number of exchange events per cell per hour. Cell segmentation and morphological analyses were performed using Cellpose. All quantitative image analyses were executed using `MATLAB` (version R2023a) or `ImageJ` (version 2.1.0/1.53c).

### Glucose and Glutamine Tracing

MDCK cells were seeded at 30,000 cells/cm^2^ in 6-well plates, ensuring that all samples for each time point were harvested simultaneously. For MPC inhibition, cells were treated with 2 *µ*M UK5099 for 80 hours. Tracing was performed by culturing cells for 24 hours in either glucose-free DMEM (Gibco, 11966-025) or glutamine-free DMEM (Gibco, 11960-044), supplemented with [U-^13^C]glucose (Cambridge Isotope Laboratories, CLM-1396-PK) or [U-^13^C]glutamine, respectively.

Before metabolite extraction, medium was aspirated and cells were washed with cold 150 mM ammonium acetate (pH 7.3). Metabolites were extracted in 500 *µ*L of cold 80% methanol, and cells were detached using a cell scraper. Norvaline (10 *µ*L of 1 mM; Sigma) was added as an internal standard. Samples were vortexed briefly and centrifuged at 17,000 *g* for 5 minutes at 1 °C. The supernatant (400 *µ*L) was transferred to glass vials (Fisher Scientific) and evaporated to dryness (EZ-2 Elite evaporator, Genevac). Dried extracts were resuspended in 300 *µ*L of 50% acetonitrile:water, centrifuged for 15 minutes at 4 °C, and 5 *µ*L of the clarified extract was injected onto a Luna NH_2_ 100 Å, 3 *µ*m, 150 × 2.0 mm column (Phenomenex). Separation was performed on a Vanquish Flex UHPLC (Thermo Scientific) using mobile phases A (5 mM NH_4_AcO, pH 9.9) and B (acetonitrile) at 200 *µ*L/min. The gradient was 15%–95% A over 18 minutes, followed by 7 minutes at 95% A and re-equilibration to 15% A. Metabolites were analyzed on a Q Exactive mass spectrometer (Thermo Scientific) operated in polarity-switching full-scan mode (70–975 m/z, 70,000 resolution).

Metabolite quantification was performed in Maven (v8.1.27.11) using accurate mass (*<*5 ppm) and verified retention times. Values were normalized to cell number or protein content. Cell counts were determined from sacrificial wells using a Countess II automated cell counter and trypan blue exclusion (average of two replicates). Relative metabolite abundances were calculated as the sum of all isotopologue intensities, corrected for natural ^13^C abundance using AccuCor. Data analysis was performed in R using custom scripts.

### Seahorse Assay

Oxygen consumption rate (OCR) and extracellular acidification rate (ECAR) were measured using a Seahorse XFe24 Extracellular Flux Analyzer (Agilent Technologies). MDCK cells were seeded at 57,000 cells per well in Seahorse XF24 plates. On the day of the assay, cells were washed and incubated in unbuffered DMEM assay medium supplemented with 8 mM glucose, 2 mM glutamine, 1 mM sodium pyruvate, and 5 mM HEPES (pH 7.4). Mitochondrial stress tests were performed by sequential injection of 2 *µ*M oligomycin (Port A), 0.75 *µ*M FCCP (Port B), 1.35 *µ*M FCCP (Port C), and a combination of 1 *µ*M rotenone and 2 *µ*M antimycin A (Port D). OCR and ECAR were recorded simultaneously throughout the assay. Rate measurements were normalized to cell number, determined by quantifying Hoechst-stained nuclei (10 *µ*g/mL) using an Operetta High-Content Imaging System (PerkinElmer). ATP production rates were calculated and data were analyzed using previously established methods ^59,60^.

### Endocytosis and Macropinocytosis Assays

Cells were seeded in 4-chamber Ibidi *µ*-slides at a density of 30,000 cells/cm^2^. To inhibit mitochondrial pyruvate uptake, cells were pre-treated with 5 *µ*M UK5099 for three days. To inhibit macropinocytosis, cells were treated with 5 *µ*M EIPA (MedChemExpress, HY-101840) for two hours prior to further analysis. For the endocytosis assay, following EIPA treatment, the cells were fixed and immunostained for the early endosome marker EEA1 using the previously mentioned immunostaining protocol. For the macropinocytosis assay, we adapted the protocol established by Le et al.^61^. Following EIPA treatment, the cells were put on ice and washed three times with cold PBS containing calcium and magnesium (Fisher Scientific, 14040-133). The cells were then incubated with 0.1mg/mL FITC-Dextran 70 (Santa Cruz Biotechnology, sc-263323) in serum-free MEM-*α* medium for 15 minutes at 37 °C. To block further internalization, the cells were immediately put back on ice, washed three times with cold PBS+Ca/Mg to remove residual dextran. Cells were fixed with 4% paraformaldehyde (Thermo Scientific, 043368.9M) for 10 minutes at room temperature and stained with DAPI for 15 minutes. Confocal imaging was performed on NL5+ with Zeiss AXIO Observer Z1 microscope using a 63 × oil/1.40 NA objective. Fluorescence intensities for EEA1 and dextran-FITC were quantified with ImageJ/Fiji and normalized to the control condition.

### Data Analysis and Statistics

All statistical analyses and data visualization were performed using GraphPad Prism (version 10) and `MATLAB` (version R2023a). Data are presented as mean ± standard deviation (SD). Correlative analyses included calculation of Pearson and Spearman correlation coefficients, along with corresponding *p*-values, false discovery rates (FDR), and confidence intervals (CI), as summarized in Table S1. FDR values were estimated using Monte Carlo simulations by random permutation of *X*–*Y* data pairs. Statistical significance was defined as follows: *p >* 0.05, not significant (ns); *p* ≤ 0.05, *; *p* ≤ 0.01, **; *p* ≤ 0.001, ***; and *p* ≤ 0.0001, ****.

## Supporting information

MPC inhibition enhances collective cell motility.

Representative T1 transition in control cells.

Representative T1 transition in UK5099-treated cells.

F-actin dynamics and baseline membrane ruffling in control cells.

Enhanced membrane ruffling following UK5099 treatment.

## Acknowledgements

We thank Fridtjof Brauns for insightful discussions. N.Y.C.L. was supported by NSF (CBET-2244760, CMMI-2029454, DBI-2325121) and NIH NIGMS (R35GM146735). J.K.H. was supported by NIH NIDCR (R01DE030471). D.B. was supported by the NSF (DMR-2046683), the NIH NIGMS (R35GM15049), the NSF (PHY-2019745) and the Alfred P. Sloan Foundation.

## Author contributions statement

A.B., J.K.H., and N.Y.C.L. conceived the project. A.B., Z.L., and J.D. performed the experiments. A.B., Z.L., and J.C. performed the cell line generation. A.B., Z.L., W.Y., D.B., A.G., J.K.H., and N.Y.C.L analyzed data. A.B., Z.L., D.B., A.G., J.K.H, and N.Y.C.L. wrote and reviewed the manuscript.

