## Extended figures for "Cell jamming transition is regulated by mitochondrial pyruvate transport and endocytosis"

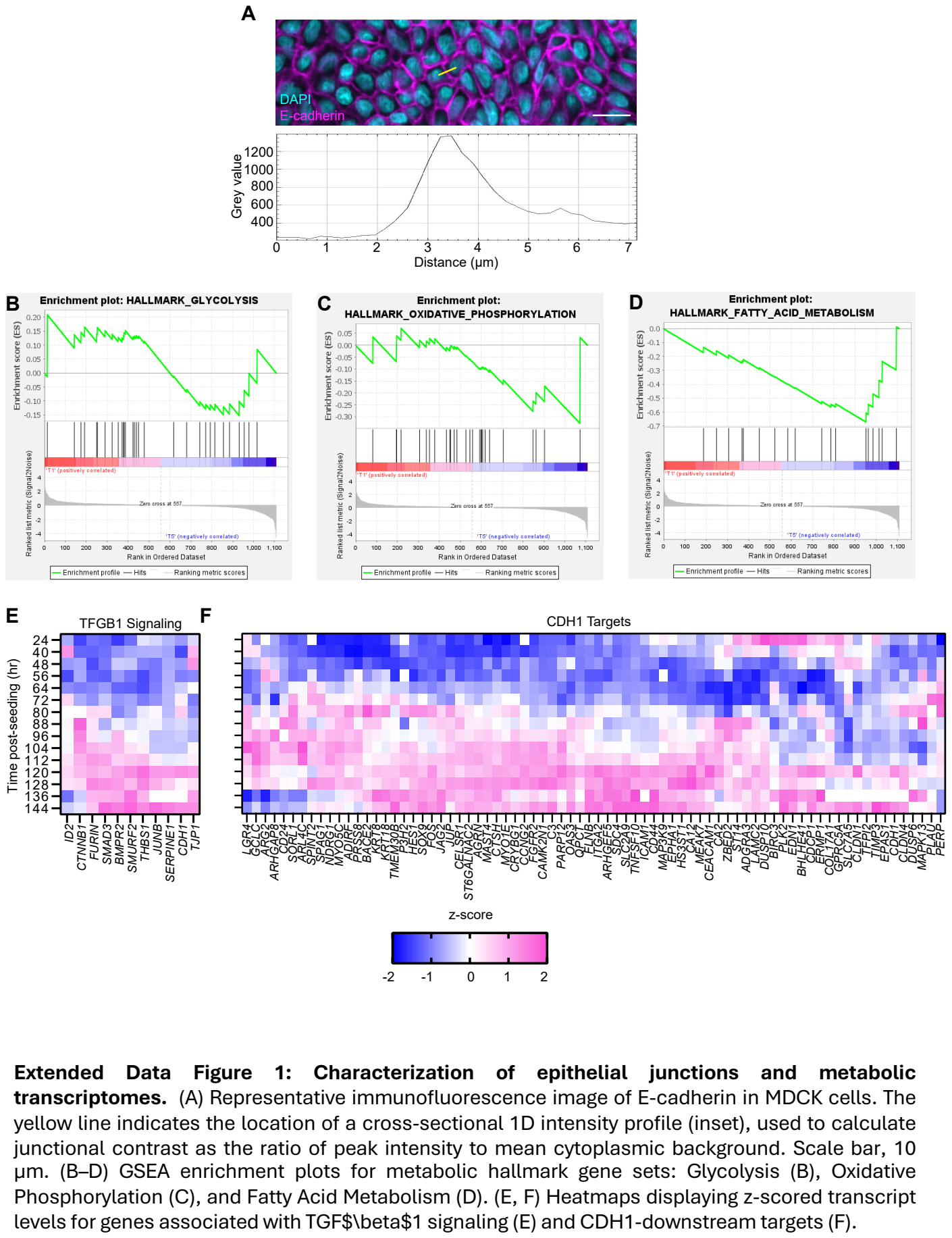

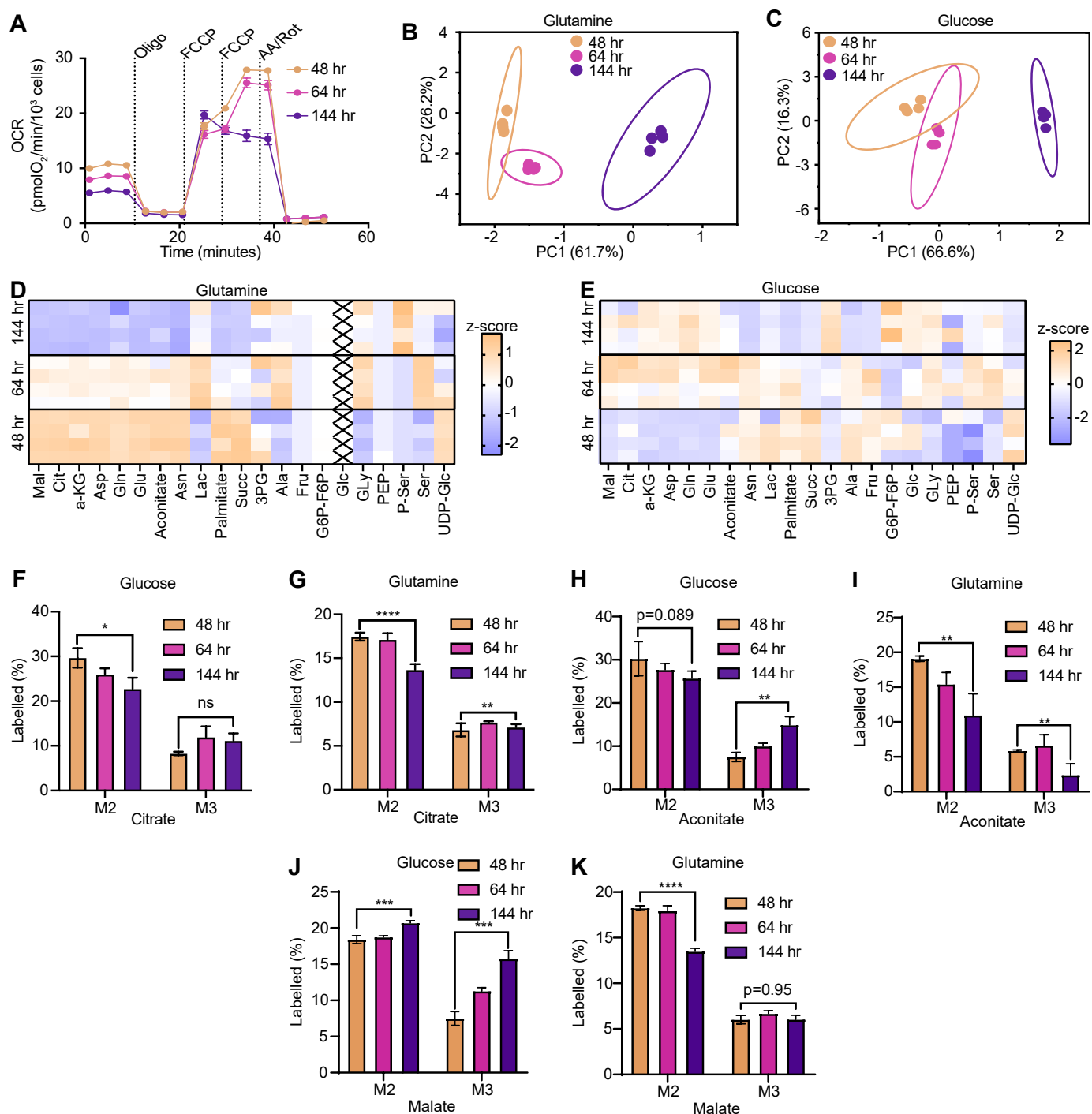

**Extended Data Figure 2: Metabolic remodeling during epithelial crowding.** (A) Oxygen consumption rate (OCR) kinetic profiles during a mitochondrial stress test (Seahorse XF) across the indicated time points. OCR values are normalized to cell density. Vertical dashed lines indicate the sequential injection of oligomycin (Oligo), FCCP, and antimycin A/rotenone (AA/Rot). (B, C) Principal Component Analysis of fractional contributions from (B) [U-<sup>13</sup>C] glutamine and (C) [U-<sup>13</sup>C] glucose tracing. Distinct clustering by time point indicates temporal shifts in substrate utilization. (D, E) Heatmaps of z-scored fractional contributions for TCA cycle intermediates derived from glutamine (D) and glucose (E). (F–K) Mass isotopologue distributions (MIDs) for citrate (F, G), aconitate (H, I), and malate (J, K), highlighting M+2 and M+3 enrichments. Results are shown for glucose tracing (F, H, J) and glutamine tracing (G, I, K). ns, \*, \*\*, \*\*\* and \*\*\*\* correspond to p-values > 0.05, ≤ 0.05, ≤ 0.01, ≤ 0.001, and ≤ 0.0001, respectively.

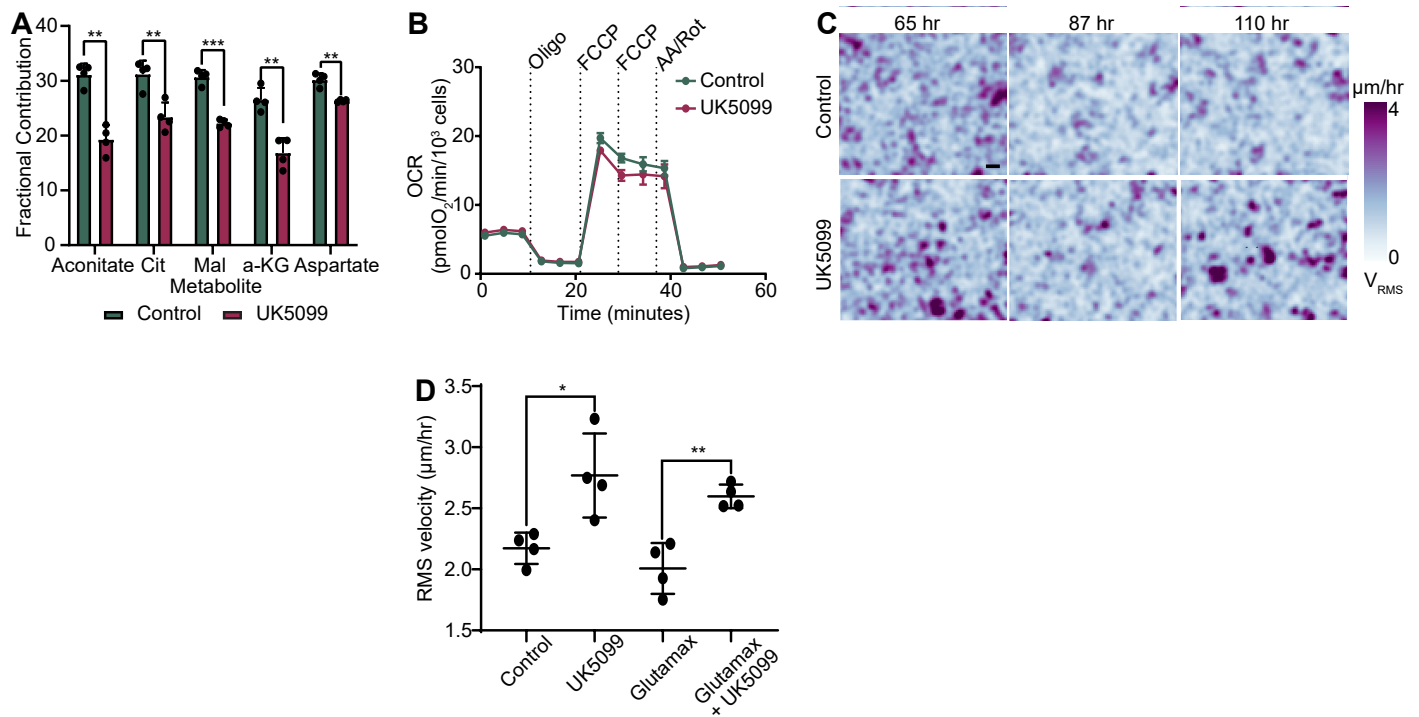

**Extended Data Figure 3: MPC inhibition via UK5099 promotes cell motility during epithelial crowding.** (A) [U-<sup>13</sup>C] glucose tracing analysis of TCA cycle intermediates following treatment with the MPC inhibitor, UK5099. Reduced fractional contributions confirm successful inhibition of glucose-derived pyruvate entry into the TCA cycle. (B) Oxygen consumption rate (OCR) kinetic profiles (Seahorse XF) showing negligible impact of UK5099 on overall mitochondrial respiration. (C) Representative heatmaps of root-mean-square velocity ($V_{rms}$) at indicated time points for control and UK5099-treated monolayers. Scale bar, 10 $\mu\text{m}$. (D) Comparison of motility phenotypes in media supplemented with L-glutamine versus GlutaMAX to examine potential glutamine degradation effects. ns, \*, \*\*, \*\*\* and \*\*\*\* correspond to p-values $> 0.05$, $\leq 0.05$, $\leq 0.01$, $\leq 0.001$, and $\leq 0.0001$, respectively.

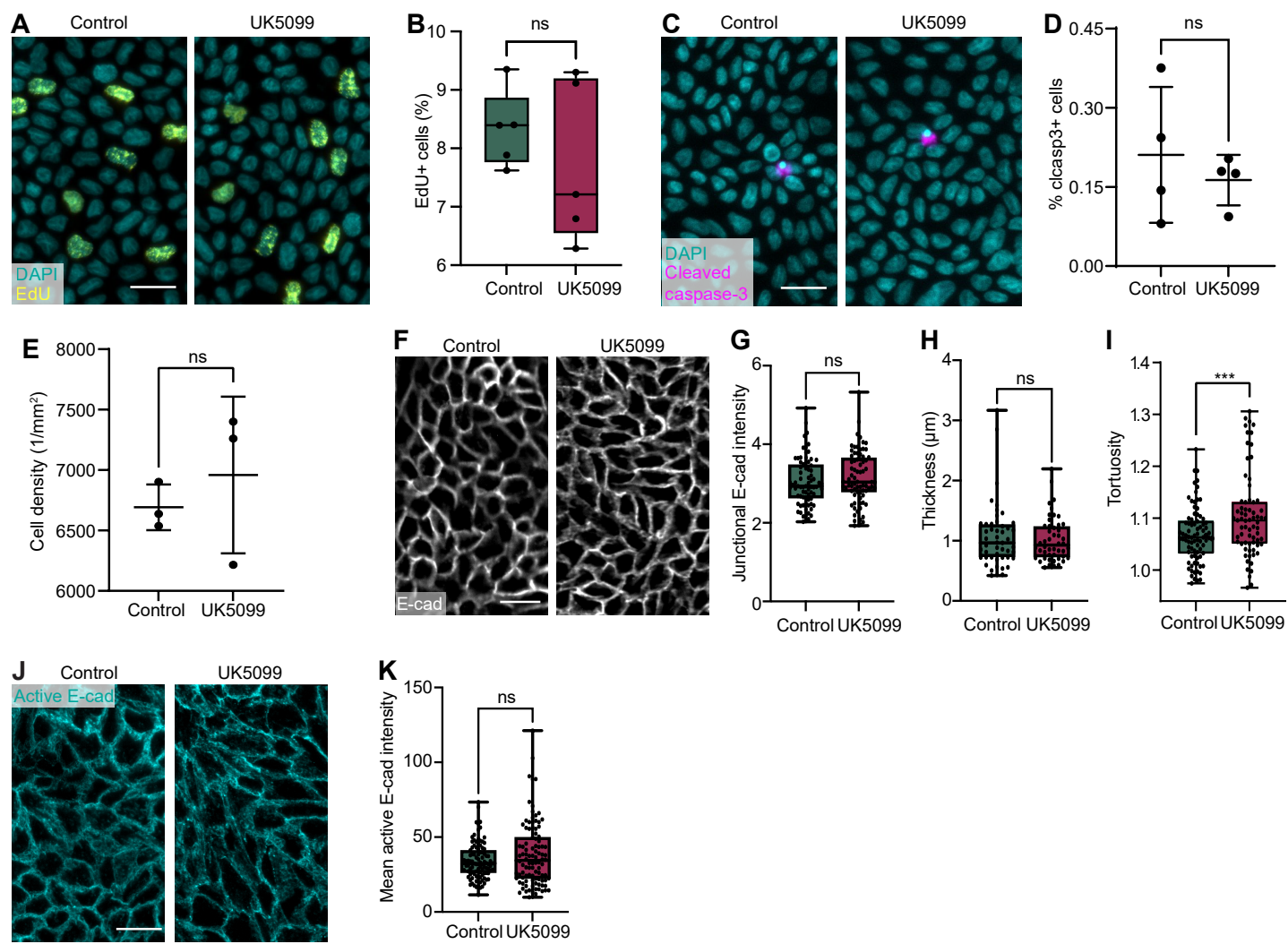

**Extended Data Figure 4: Impact of MPC inhibition on cell proliferation, apoptosis, and junctional integrity.** (A, B) Representative immunofluorescence images of EdU incorporation (A) and subsequent quantification (B), showing comparable proliferation rates between control and UK5099-treated cells. (C, D) Representative images of cleaved caspase-3 (clCASP3) staining (C) and quantification of clCASP3+ cells (D), indicating no significant induction of apoptosis upon UK5099 treatment. (E) Total cell density measurements across the indicated conditions. (F–I) Analysis of E-cadherin junctional morphology. Representative immunofluorescence images of E-cadherin (F) and quantification of junctional intensity (G), junctional thickness (H), and tortuosity (I). Tortuosity is defined as the ratio of the curvilinear junctional length to the Euclidean end-to-end distance. (J, K) Immunofluorescence of homophilically bound (active) E-cadherin (J) and corresponding quantification of mean fluorescence intensity (K). ns, \*, \*\*, \*\*\* and \*\*\*\* correspond to p-values $> 0.05$, $\leq 0.05$, $\leq 0.01$, $\leq 0.001$, and $\leq 0.0001$, respectively. Scale bars, 50 $\mu\text{m}$.

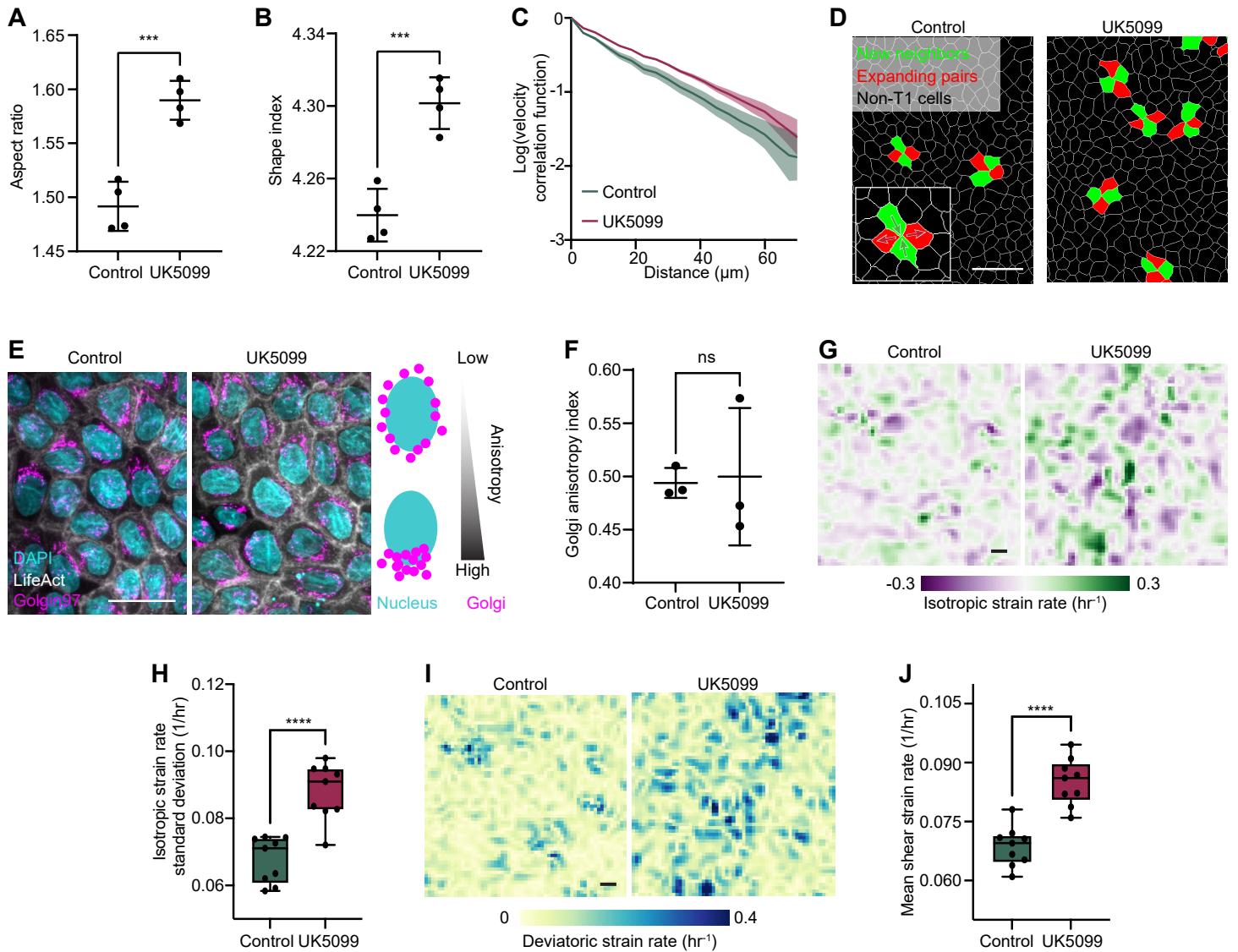

**Extended Data Figure 5: Morphological and kinematic characterization of MPC-inhibited monolayers.** (A, B) Quantification of cell (A) aspect ratio and (B) shape index in control and UK5099-treated monolayers. (C) Spatial velocity-velocity correlation functions as a function of intercellular distance. UK5099 treatment results in an increased correlation length, characterized by a slower spatial decay of the correlation metric. (D) Representative snapshot from TissueAnalyzer illustrating the identification and tracking of T1 transition events. (E, F) Analysis of cell front-rear polarity via Golgin-97 immunofluorescence. (E) Representative images; the inset illustrates the method for calculating the intensity angular anisotropy index. (F) Quantification of the anisotropy index shows that MPC inhibition does not significantly alter global cell polarization. (G, H) Representative heatmaps of the isotropic strain rate derived from the velocity field (G) and corresponding quantification (H), showing enhanced volumetric expansion/contraction rates upon UK5099 treatment. (I, J) Representative heatmaps of the deviatoric strain rate (I) and quantification (J), indicating that MPC inhibition increases shear-like deformations within the monolayer. ns, \*, \*\*, \*\*\* and \*\*\*\* correspond to p-values $> 0.05$, $\leq 0.05$, $\leq 0.01$, $\leq 0.001$, and $\leq 0.0001$, respectively. Scale bars, 50 $\mu\text{m}$.

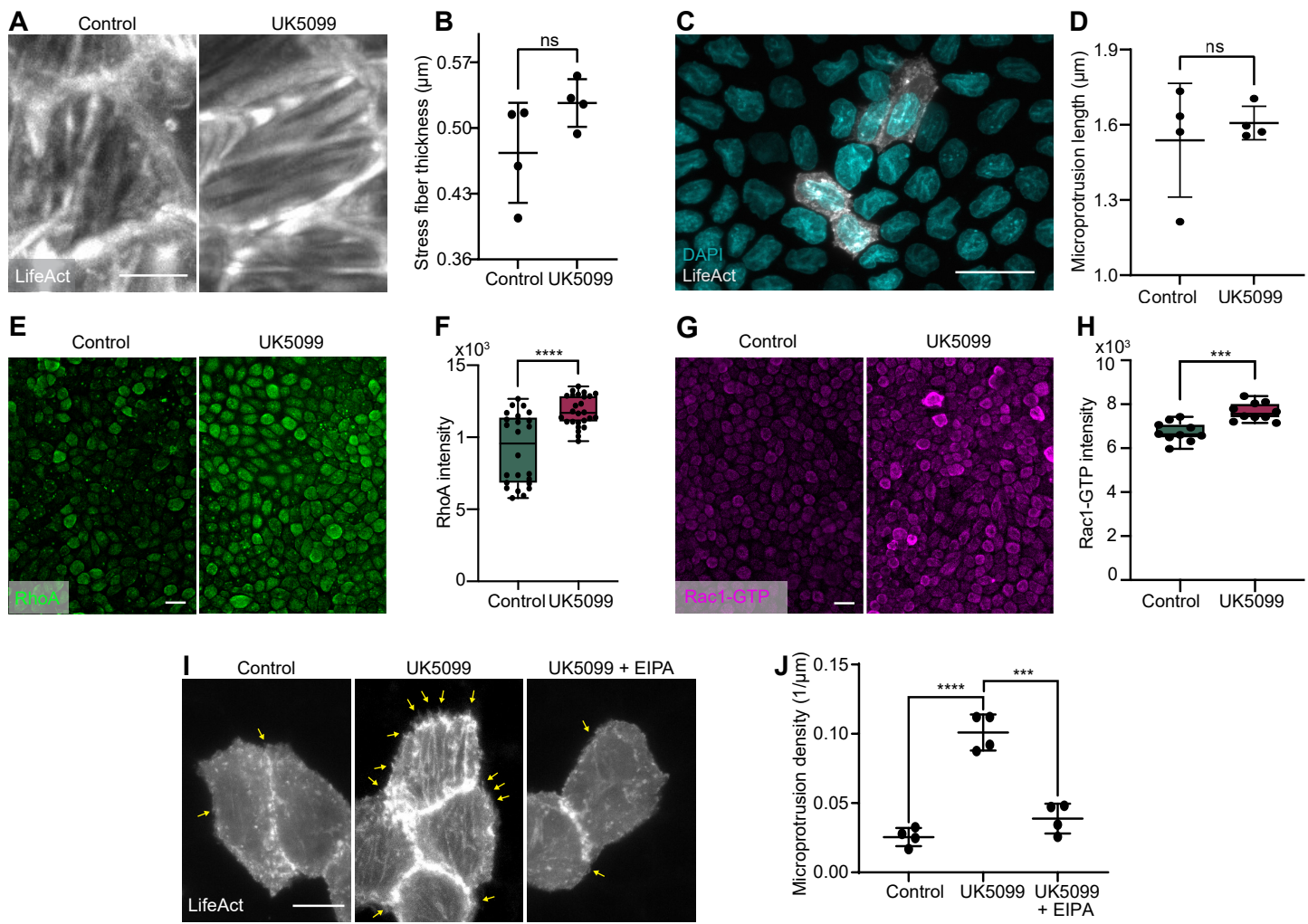

**Extended Data Figure 6: Cytoskeletal dynamics and Rho-family GTPase activity upon MPC inhibition.** (A, B) Representative fluorescence images of LifeAct-GFP (A) and quantification of stress fiber thickness (B), showing preserved contractile fiber morphology following UK5099 treatment. (C) Representative field of view illustrating the cell density and distribution for sparse labeling experiments. (D) Quantification of microprotrusion length, indicating that MPC inhibition does not significantly alter protrusion length. (E, F) Immunofluorescence images of total RhoA (E) and corresponding intensity quantification (F), showing an upregulation of RhoA expression in UK5099-treated cells. (G, H) Immunofluorescence images of Rac1-GTP (G) and corresponding quantification (H), demonstrating increased levels of the active GTP-bound form of Rac1 upon MPC inhibition. (I, J) Representative fluorescence images of LifeAct-GFP (I) and quantification of microprotrusion linear density (J). UK5099-induced microprotrusions (yellow arrows) are significantly suppressed following treatment with the $\text{Na}^+/\text{H}^+$ exchanger inhibitor EIPA. ns, \*, \*\*, \*\*\* and \*\*\*\* correspond to p-values $> 0.05$, $\leq 0.05$, $\leq 0.01$, $\leq 0.001$, and $\leq 0.0001$, respectively. Scale bars, 50 $\mu\text{m}$.

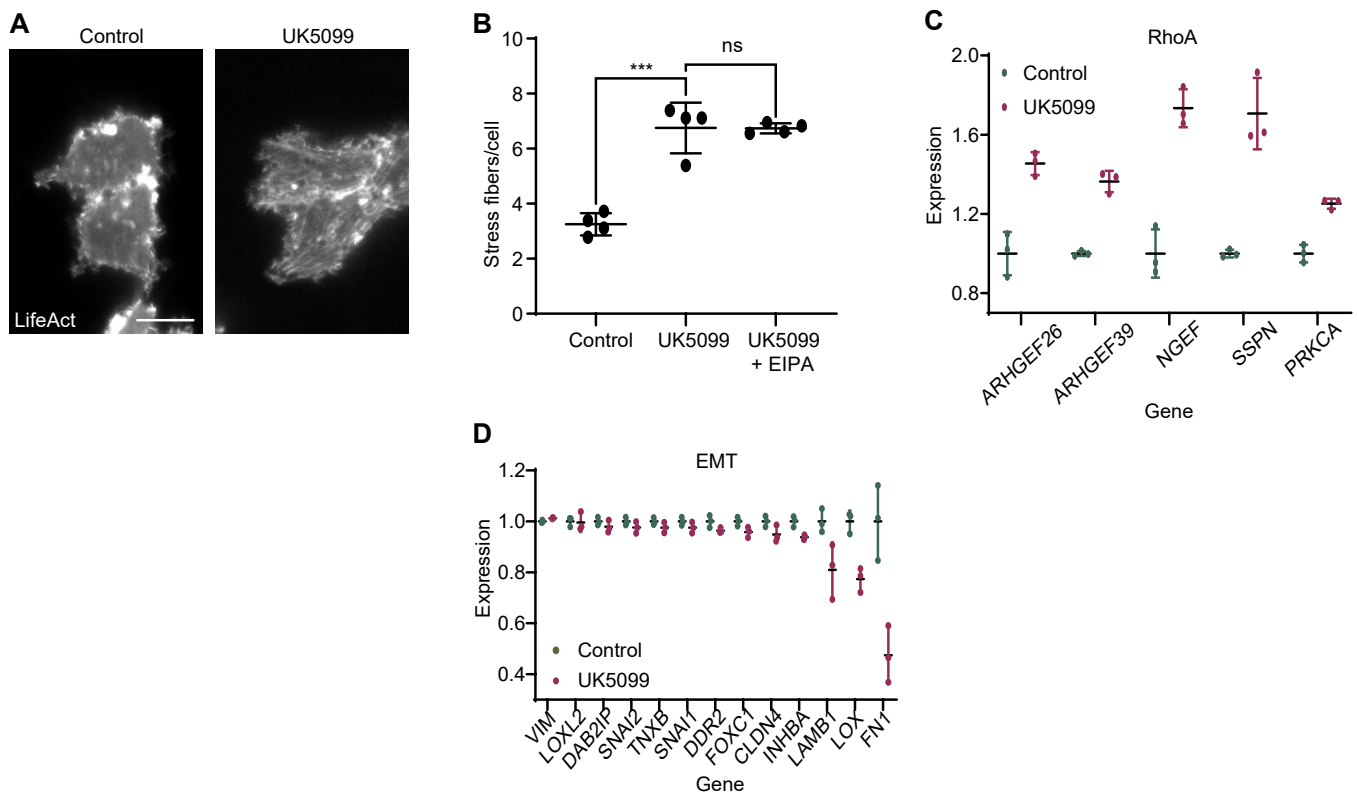

**Extended Data Figure 7: Impact of EIPA and MPC inhibition on cytoskeletal architecture and transcriptomic profiles.** (A, B) Representative fluorescence images of LifeAct-GFP (A) and quantification of stress fiber density (B) in control and EIPA-treated monolayers. $\text{Na}^+/\text{H}^+$ exchanger exchange inhibition via EIPA does not significantly alter stress fiber formation. ns, \*, \*\*, \*\*\* and \*\*\*\* correspond to p-values $> 0.05$, $\leq 0.05$, $\leq 0.01$, $\leq 0.001$, and $\leq 0.0001$, respectively. Scale bar, 10 $\mu\text{m}$. (C) Differential gene expression analysis via RNA-seq illustrating the upregulation of transcripts associated with RhoA signaling pathways. (D) Transcript levels of core EMT markers from RNA-seq data. No significant induction of a mesenchymal signature was observed, with specific markers (LAMB1, LOX, and FN1) showing modest downregulation upon UK5099 treatment. These results suggest that MPC inhibition-mediated motility occurs independently of a transition to a mesenchymal cell fate.
